## Supplementary Information for "Enhancement of Enzymatic Activity by Biomolecular Condensates through pH Buffering"

| **Protein** | **Sequence** |
|---|---|
| BTL2 | MGSSHHHHHHSSGLVPRGSHMMKGCRVMVVLLGLWFVFGLSVPGGRTEAASPRANDAPIVLLH<br>GFTGWGREEMLGFKYWGGVRGDIEQWLNDNGYRTYTLAVGPLSSNWDRACEAYAQLVGGTVD<br>YGAAHAAKHGHARFGRTYPGLLPELKRGGRVHIIAHSQGGQTARMLVSLLENGSQEEREYAKAHN<br>VSLSPLFEGGHHFVLSVTTIATPHDGTTLVNMVDFTDRFFDLQKAVLKAAAVASNVPYTSQVYDFKL<br>DQWGLRRQPGESFDHYFERLKRSPVWTSTDTARYDLSIPGAEKLNQWVQASPNTYYLSFSTERTH<br>RGALTGNYYPELGMNAFSAVVCAPFLGSYRNEALGIDDRWLENDGIVNTVSMNGPKRGSSDRIVP<br>YDGTLKKGVWNDMGTCNVDHLEVIGVDPNPSFDIRAFYLRLAEQLASLRP |
| Laf1-BTL2-Laf1 | MGSSHHHHHHSSGLVPRGSHMESNQSNNGGSGNAALNRGGRYVPPHLRGGDGGAAAAASAG<br>GDDRRGGAGGGGYRRGGGNSGGGGGGGYDRGYNDNRDDRDNRGGSGGYGRDRNYEDRGYN<br>GGGGGGGNRGYNNNRGGGGGGYNRQDRGDGGSSNFSRGGYNNRDEGSDNRGSGRSYNNDR<br>RDNGGDGASPRANDAPIVLLHGFTGWGREEMLGFKYWGGVRGDIEQWLNDNGYRTYTLAVGPL<br>SSNWDRACEAYAQLVGGTVDYGAAHAAKHGHARFGRTYPGLLPELKRGGRVHIIAHSQGGQTAR<br>MLVSLLENGSQEEREYAKAHNVSLSPLFEGGHHFVLSVTTIATPHDGTTLVNMVDFTDRFFDLQKA<br>VLKAAAVASNVPYTSQVYDFKLDQWGLRRQPGESFDHYFERLKRSPVWTSTDTARYDLSIPGAEKL<br>NQWVQASPNTYYLSFSTERTHRGALTGNYYPELGMNAFSAVVCAPFLGSYRNEALGIDDRWLEND<br>GIVNTVSMNGPKRGSSDRIVPYDGTLKKGVWNDMGTCNVDHLEVIGVDPNPSFDIRAFYLRLAEQ<br>LASLRPMESNQSNNGGSGNAALNRGGRYVPPHLRGGDGGAAAAASAGGDDRRGGAGGGGYRR<br>GGGNSGGGGGGGYDRGYNDNRDDRDNRGGSGGYGRDRNYEDRGYNGGGGGGGNRGYNNN<br>RGGGGGGYNRQDRGDGGSSNFSRGGYNNRDEGSDNRGSGRSYNNDRRDNGGDG |
| Laf1 | MGSSHHHHHHSSGLVPRGSHMESNQSNNGGSGNAALNRGGRYVPPHLRGGDGGAAAAASAG<br>GDDRRGGAGGGGYRRGGGNSGGGGGGGYDRGYNDNRDDRDNRGGSGGYGRDRNYEDRGYN<br>GGGGGGGNRGYNNNRGGGGGGYNRQDRGDGGSSNFSRGGYNNRDEGSDNRGSGRSYNNDR<br>RDNGGDG |

| | |
|---|---|
| DDX4-BTL2-DDX4 | MGSSHHHHHHSSGLVPRGSHMGDEDWEAEINPHMSSYVPIFEKDRYSGENGDNFNRTPASSSE<br>MDDGPSRRDHFMKSGFASGRNFGNRDAGECNKRDNTSTMGGFGVGKSFGNRGFSNSRFEDGD<br>SSGFWRESSNDCEDNPTRNRGFSKRGGYRDGNNSEASGPYRRGGRGSFRGCRGGFGLGSPNNDL<br>DPDECMQRTGGLFGSRRPVLSGTGNGDTSQSRSGSGSERGGYKGLNEEVITGSGKNSWKSEAEG<br>GESASPRANDAPIVLLHGFTGWGREEMLGFKYWGGVRGDIEQWLNDNGYRTYTLAVGPLSSNW<br>DRACEAYAQLVGGTVDYGAAHAAKHGHARFGRTYPGLLPELKRGGRVHIIAHSQGGQTARMLVS<br>LLENGSQEEREYAKAHNVSLSPLFEGGHHFVLSVTTIATPHDGTTLVNMVDFTDRFFDLQKAVLKA<br>AAVASNVPYTSQVYDFKLDQWGLRRQPGESFDHYFERLKRSPVWTSTDTARYDLSIPGAEKLNQW<br>VQASPNTYYLSFSTERTHRGALTGNYYPELGMNAFSAVVCAPFLGSYRNEALGIDDRWLENDGIVN<br>TVSMNGPKRGSSDRIVPYDGTLKKGVWNDMGTCNVDHLEVIGVDPNPSFDIRAFYLRLAEQLASL<br>RPGDEDWEAEINPHMSSYVPIFEKDRYSGENGDNFNRTPASSSEMDDGPSRRDHFMKSGFASGR<br>NFGNRDAGECNKRDNTSTMGGFGVGKSFGNRGFSNSRFEDGDSSGFWRESSNDCEDNPTRNRG<br>FSKRGGYRDGNNSEASGPYRRGGRGSFRGCRGGFGLGSPNNDLDPDECMQ |
| AAOx | MGSSHHHHHHSSGLVPRGSHMATNLPTADFDYVVVGAGNAGNVVAARLTEDPDVSVLVLEAGV<br>SDENVLGAEAPLLAPGLVPNSIFDWNYTTTAQAGYNGRSIAYPRGRMLGGSSSVHYMVMMRGST<br>EDFDRYAAVTGDEGWNWDNIQQFVRKNEMVVPPADNHNTSGEFIPAVHGTNGSVSISLPGFPTP<br>LDDRVLATTQEQSEEFFFNPDMGTGHPLGISWSIASVGNGQRSSSSTAYLRPAQSRPNLSVLINAQ<br>VTKLVNSGTTNGLPAFRCVEYAEQEGAPTTTVCAKKEVVLSAGSVGTPILLQLSGIGDENDLSSVGID<br>TIVNNPSVGRNLSDHLLLPAAFFVNSNQTFDNIFRDSSEFNVDLDQWTNTRTGPLTALIANHLAWL<br>RLPSNSSIFQTFPDPAAGPNSAHWETIFSNQWFHPAIPRPDTGSFMSVTNALISPVARGDIKLATSN<br>PFDKPLINPQYLSTEFDIFTMIQAVKSNLRFLSGQAWADFVIRPFDPRLRDPTDDAAIESYIRDNANTI<br>FHPVGTASMSPRGASWGVVDPDLKVKGVDGLRIVDGSILPFAPNAHTQGPIYLVGKQGADLIKAD<br>Q |

**Figure S1:** Amino acid sequences of BTL2, Laf1-BTL2-Laf1, Laf1, AAOx, and DDX4-BTL2-DDX4 proteins. The N-terminal His-Tag is highlighted in red, the Laf1 IDR in grey, the BTL2 globular protein in green, the DDX4 IDR in cyan, and the AAOx globular protein in magenta.

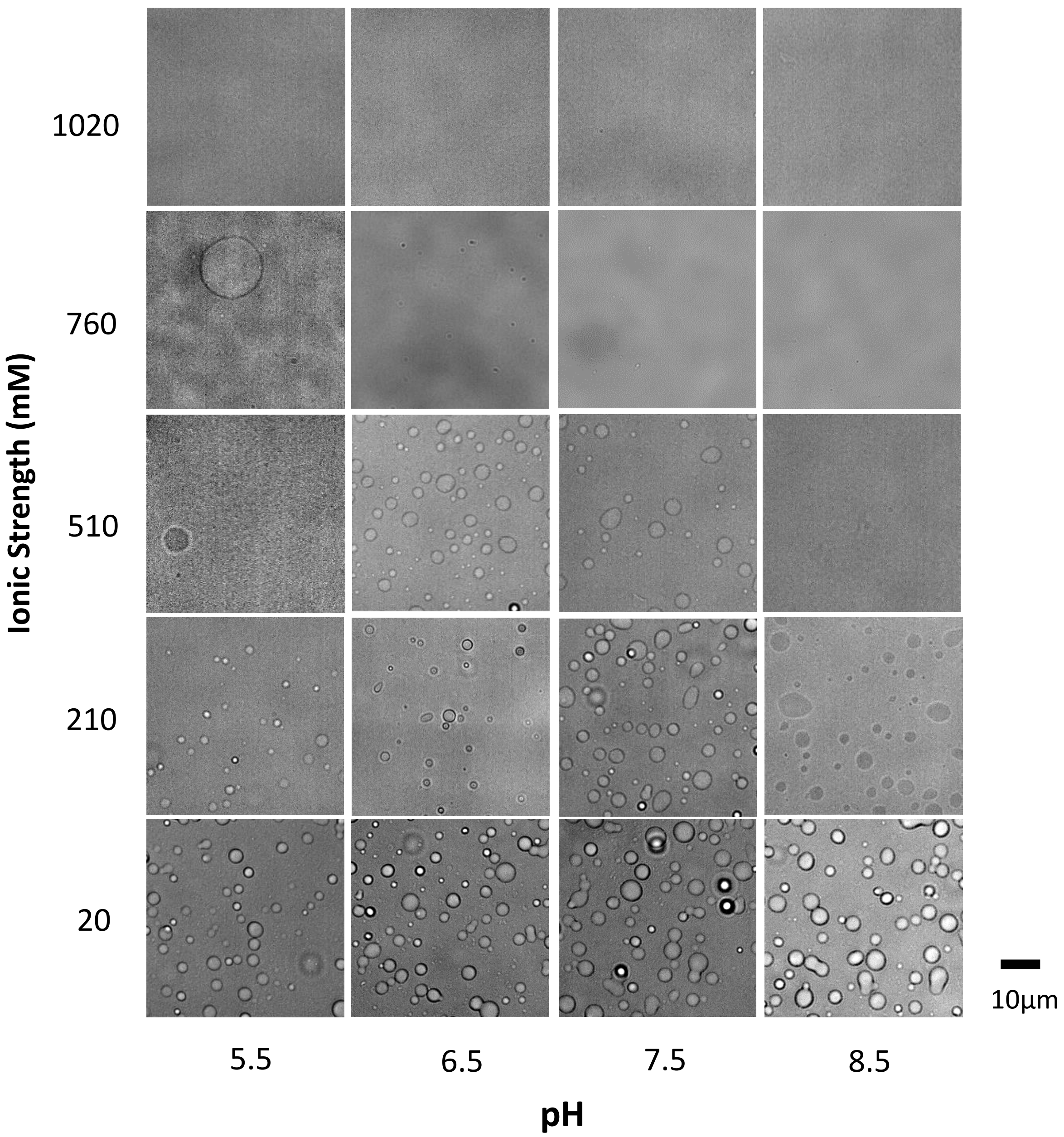


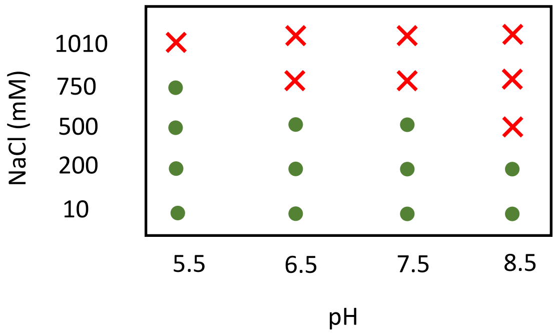


**Figure S2:** Phase diagram of 0.5 µM Laf1-BTL2-Laf1 in Tris / Bis-tris buffers at an ionic strength of 10 mM. Images were taken 10 mins after sample preparation.

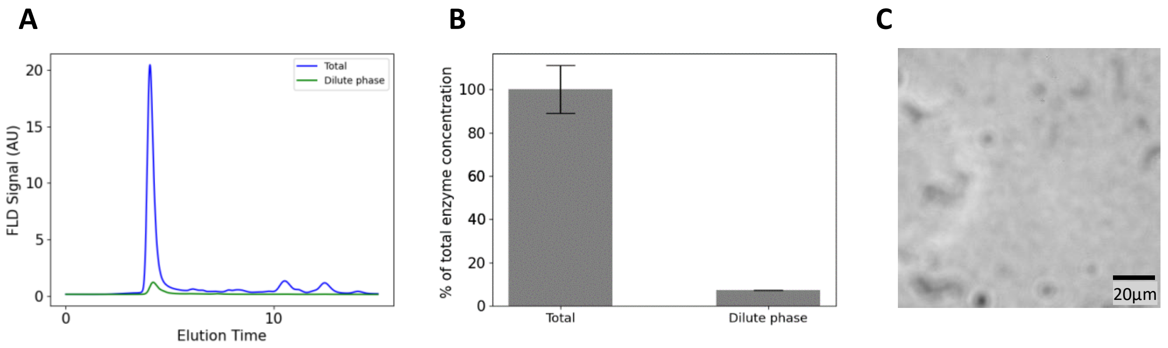


**Figure S3:** Analysis of enzyme partitioning in the dense phase via size exclusion chromatography: **A)** SEC chromatograms recorded with protein intrinsic fluorescence signal of the homogeneous Laf1-BTL2-Laf1 solution at 0.5 µM and of the dilute phase in the phase separated system (see Materials and Methods). **B)** Normalized amount of protein in the dilute phase determined by integration of main peaks corresponding to Laf1-BTL2-Laf1 in the chromatograms shown in panel **A.** **C)** Bright-field image of the dilute phase separated from the dense phase by centrifugation.

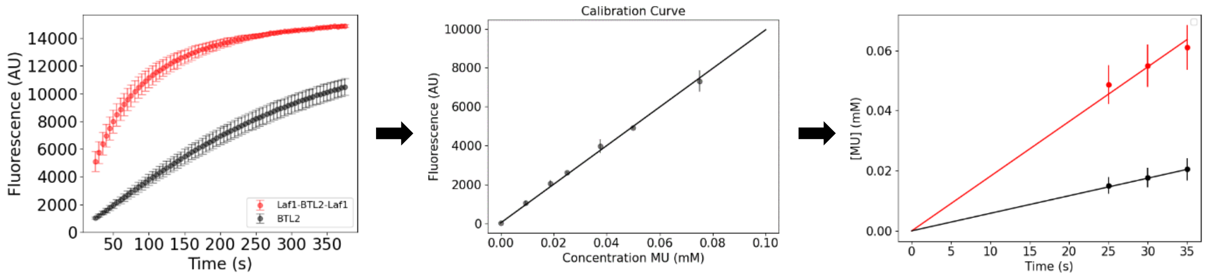


**Figure S4:** Measurement of initial rates from the complete kinetic profiles of the hydrolysis catalysed by BTL2. Linear Regressions were fitted with Graphpad Prism.

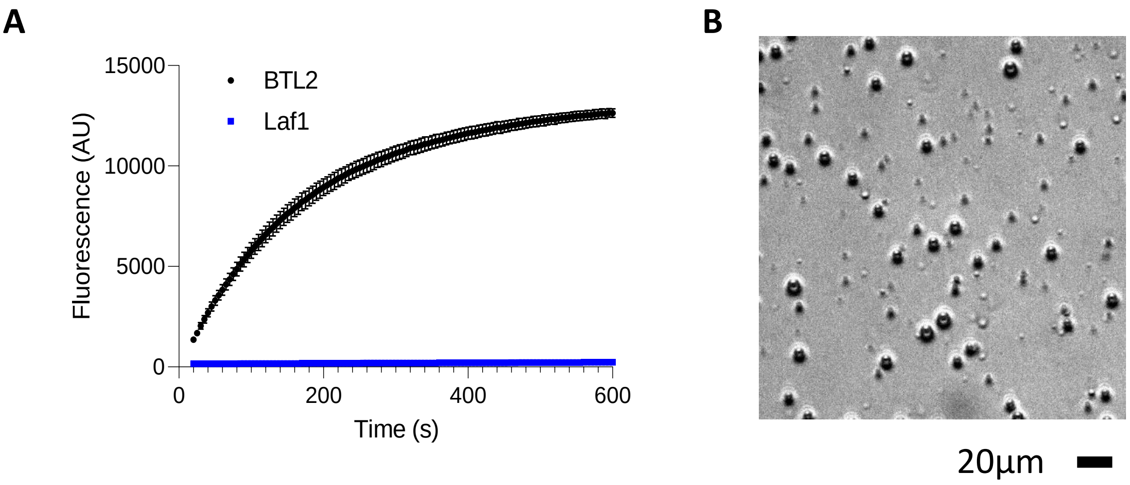


**Figure S5: A)** MUB hydrolysis over time in presence of BTL2 (black symbols, 0.5 µM protein and 0.1 mM MUB in 24 mM Tris buffer at pH 7.5 and 10 mM NaCl) and Laf1 IDR condensates (blue symbols, 10 µM protein and 0.1 mM MUB in 24 mM Tris buffer at pH 7.5 and 13 mM NaCl, 30mM Urea). **B)** Brightfield microscopy image of Laf1 IDR condensates before adding the substrate for the experiment shown in panel **A.**

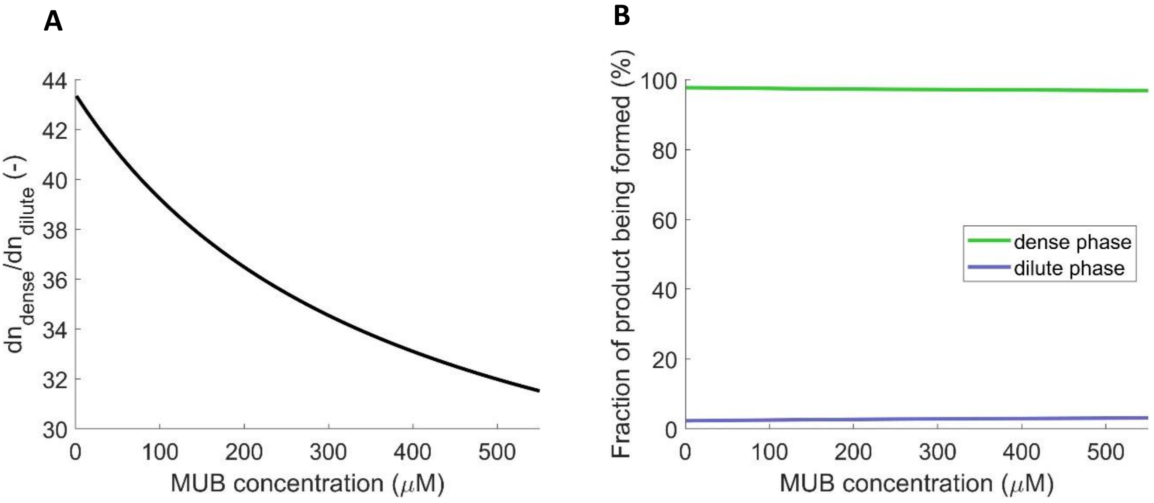


**Figure S6: A)** Ratio of product formation in the dense and dilute phases. **B)** Fraction of product formed in the dense and dilute phases, evaluated from equation (2)

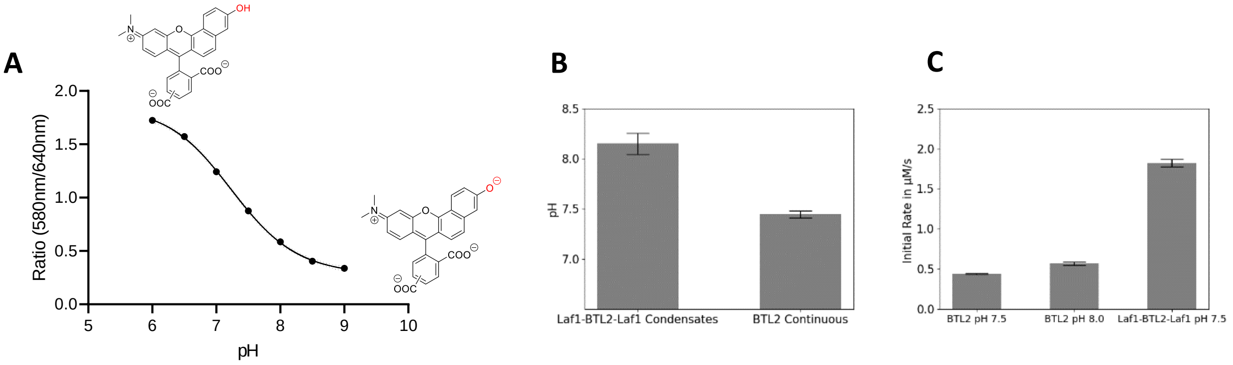


**Figure S7: A)** SNARF-1 calibration curve with associated protonation states. All buffers have 10 mM ionic strength. **B)** Apparent pH within Laf1-BTL2-Laf1 condensates compared to the dilute BTL2 phase measured with the SNARF1 assay (0.5 µM protein in 24 mM Tris Buffer at pH 7.5 and 10 mM NaCl). **C)** Initial rate of MUB hydrolysis catalyzed by BTL2 at pH 7.5 and pH 8.0 compared to that of Laf1-BTL2-Laf1 at pH 7.5 (0.5 µM protein in Tris buffer at 10 mM ionic strength and 10mM NaCl)

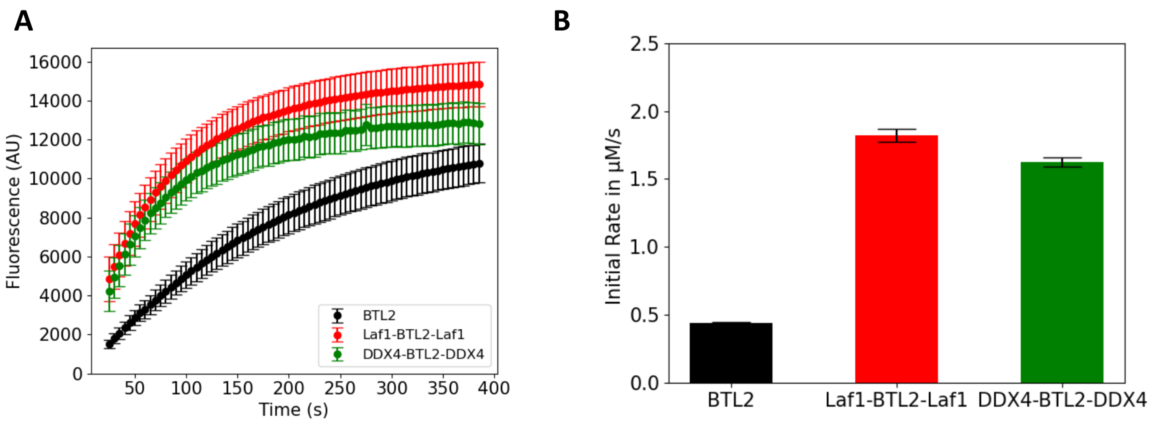


**Figure S8: A)** Kinetic curves of MUB hydrolysis catalyzed by BTL2, Laf1-BTL2-Laf1, and DDX4-BTL2-DDX4 (0.5 µM protein and 0.25mM MUB in 24 mM Tris buffer at pH 7.5, with 10 mM NaCl for Laf1-BTL2-Laf1 and BTL2, and 30 mM NaCl for DDX4-BTL2-DDX4). **B)** Initial rates extracted from the kinetic curves shown in panel **A.**

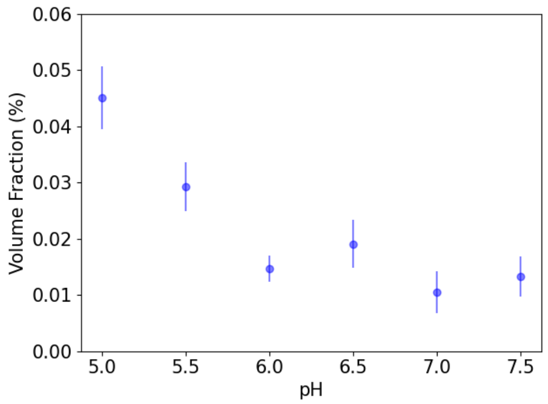


**Figure S9:** Volume fraction of the dense phase of DDX4-BTL2-DDX4 condensates at different pH values. Volume fraction measured by z-stack analysis using confocal microscopy for a 0.5µM protein sample in Tris or BisTris buffers at 10 mM ionic strength and 30 mM NaCl.

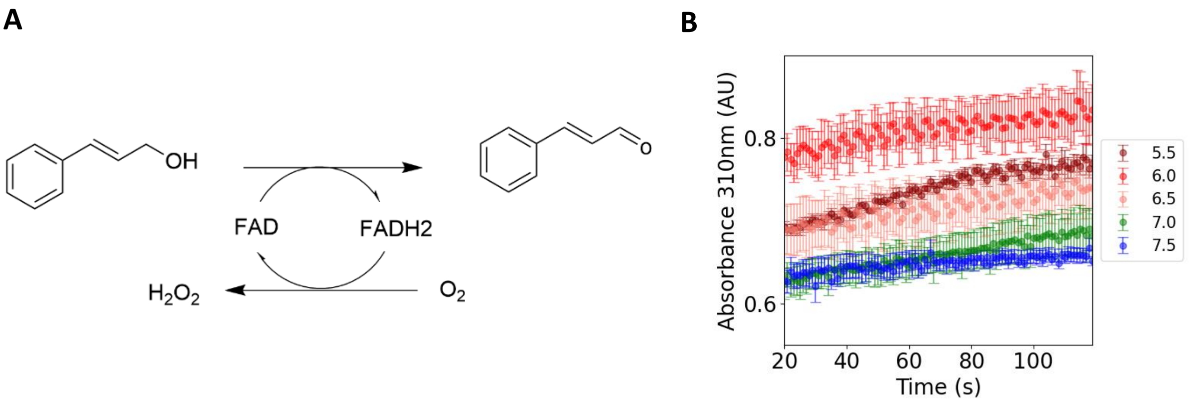


**Figure S10: A)** Scheme of the oxidation reaction of CAlc to CAld by AAOx . **B)** Representative kinetic profiles of CAlc oxidation catalyzed by AAOx measured by monitoring the absorbance of the product (310 nm) across different pH values (5 nM enzyme in Tris/ Bis-Tris buffer at 10 mM ionic strength).

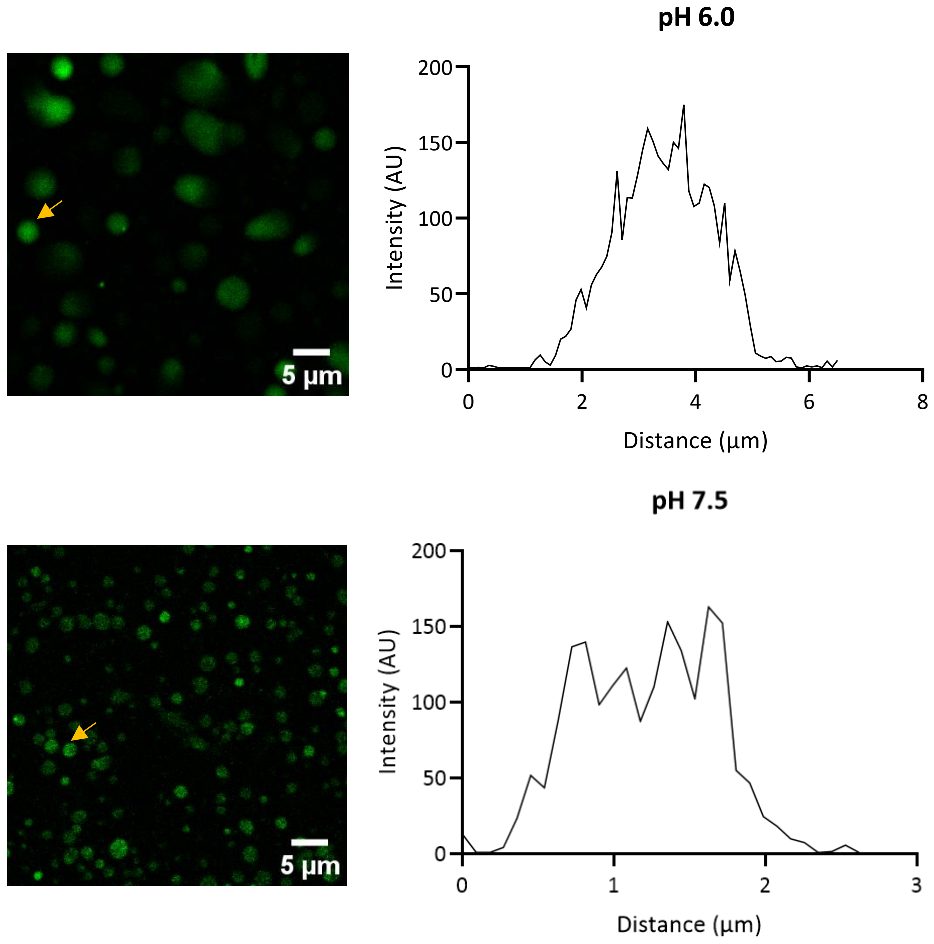


**Figure S11:** Representative intensity profiles of confocal microscopy images on the left showing the uptake of AAOx-ATTO565 into DDX4-BTL2-DDX4 condensates at pH 6.0 and pH 7.5. Data were extracted using ImageJ (see Materials and Methods). Representative analysed condensates are indicated with a yellow arrow.

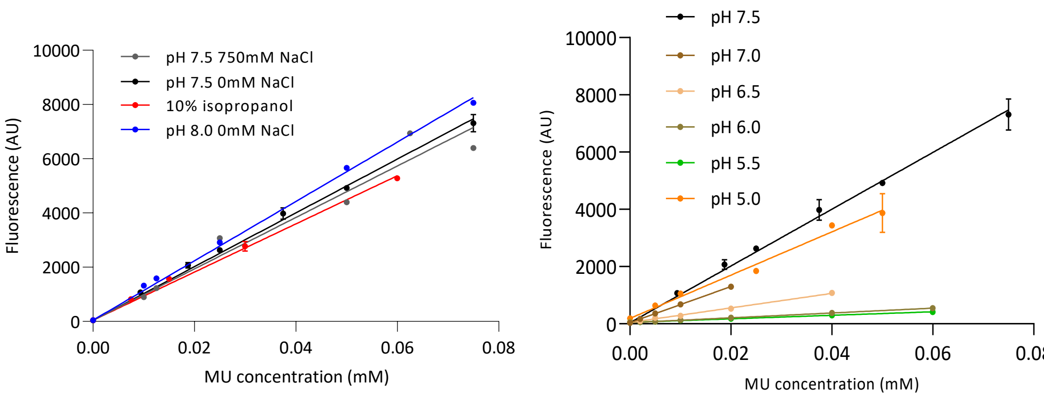


**Figure S12:** Calibration curves of fluorescence intensity versus MU concentration used in the measurement of the initial rate in different buffer conditions. For the 10% isopropanol condition, the remaining 90% consists of 24mM Tris Buffer, at pH 7.5. The buffers at different pHs are comprised of Tris or BisTris buffers with an ionic strength of 10mM.

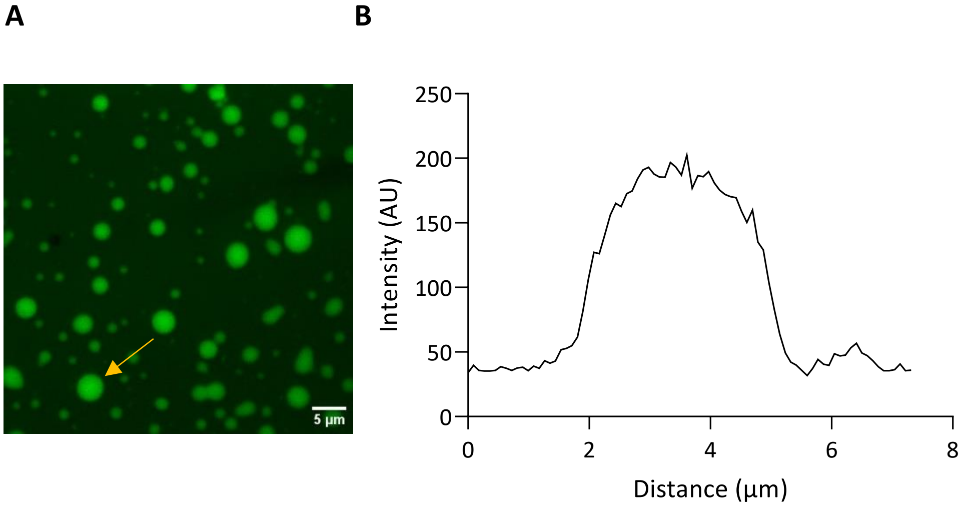


**Figure S13: A:** Fluorescence confocal microscopy image of 0.5µM Laf1-BTL2-Laf1 condensates stained with 50 µM Resorufin in 24 mM Tris buffer at pH 7.5 and 10 mM NaCl. Yellow arrow indicates the condensate corresponding to the fluorescence intensity profile shown in panel **B**.
